# Dependent variable selection in phylogenetic generalized least squares regression analysis under Pagel’s lambda model

**DOI:** 10.1101/2023.05.21.541623

**Authors:** Zheng-Lin Chen, Hong-Ji Guo, Deng-Ke Niu

## Abstract

Phylogenetic generalized least squares (PGLS) regression is widely used to examine evolutionary associations while accounting for phylogenetic non-independence, but it requires designating one trait as the dependent variable and the other as the independent variable. When causal relationships between traits are unclear, choosing which trait serves as the dependent variable becomes a practical concern. While studying the evolutionary relationship between bacterial growth rate and CRISPR-Cas content, we noticed that switching the roles of dependent and independent variables in PGLS analyses could lead to inconsistent conclusions in a substantial proportion of cases. To confirm this observation, we conducted 16,000 simulations of trait evolution along binary trees with 100 terminal nodes under different evolutionary models. PGLS regressions using Pagel’s λ model were applied to each simulation, and the results consistently showed that swapping the dependent and independent variables can lead to inconsistent outcomes. We evaluated seven potential criteria for selecting the dependent variable, including log-likelihood, AIC, R^2^, p-value, Pagel’s λ, Blomberg’s K, and the estimated λ in the Pagel’s λ model. Among these, Pagel’s λ, Blomberg’s K, and the estimated λ performed equally well and outperformed the others in selecting the dependent variable, providing a reliable basis for PGLS analyses when causal direction between traits is unclear.

## 1 INTRODUCTION

Through mutations, genetic drift, and natural selection, biological traits often co-evolve along the phylogeny of species (Felsenstein 1985; Revell & Collar 2009; Goswami *et al*., 2014; Revell *et al*., 2022). Studying correlations among such traits can provide insights into how changes in one characteristic may accompany evolutionary changes in another, offering support for or challenges to specific biological hypotheses (Pearman *et al*., 2014). Understanding these correlations contributes to reconstructing evolutionary histories and uncovering underlying biological processes (Bartomeus *et al*., 2018; Bawa *et al*., 2019; Suarez-Castro *et al*., 2020).

A common approach for assessing trait associations is through correlation analysis, using Pearson, Spearman, or Kendall coefficients. However, conventional statistical methods assume independence among observations—a condition violated in comparative biological data, where species share evolutionary history. Ignoring phylogenetic non-independence can lead to inflated or distorted correlation estimates (Revell 2010; Whitney & Garland 2010). To address this, phylogenetic comparative methods (PCMs) have been developed, explicitly accounting for the phylogenetic structure of the data (Felsenstein 1985; Grafen 1989; Lynch 1991; Garamszegi 2014; Cooper *et al*., 2016).

Among PCMs, phylogenetic generalized least squares (PGLS) has become widely used to study associations between traits. Originally proposed by Grafen (1989) and refined by Martins and Hansen (1997), Pagel (1997; 1999), and Rohlf (2001), PGLS extends ordinary least squares (OLS) regression by incorporating the expected covariance among species due to shared ancestry. Despite being fundamentally a regression method, PGLS is frequently used to examine trait correlations, likely due to the lack of simple and flexible tools for phylogenetically controlled correlation analysis. While other approaches such as phylogenetic independent contrasts (PIC) can also estimate correlations (Felsenstein 1985), PIC is limited to traits evolving under a Brownian motion (BM) model and cannot accommodate traits with more complex evolutionary dynamics.

Regression-based methods inherit OLS assumptions, including that one variable is the independent variable (predictor) and the other is the dependent variable (response), implying a causal direction (Waugh 1943). In practice, however, the causal relationships among many biological traits are often uncertain or debated for decades. For example, the relationship between GC content and environmental temperature has been discussed in multiple ways: one hypothesis suggests GC content affects DNA double-strand stability, influencing thermal tolerance, while another emphasizes that temperature-induced DNA damage may not produce symmetric changes in GC-AT content(Travnicek *et al*., 2019; Hu *et al*., 2022). Each perspective can support an alternative causal direction, and from a biological standpoint, there is often no basis to prioritize one over the other.

When applying PGLS to study the correlation between two traits with unclear causal relationships, the assignment of dependent and independent variables may not matter if swapping them does not change the outcome—i.e., the sign (positive or negative) of the regression coefficient or the significance test results (*p* < or ≥ 0.05). In practice, however, we found that using PGLS with Pagel’s λ model (Freckleton et al., 2002) on simulated data, swapping dependent and independent variables could substantially affect the results: correlations that were significant in one configuration sometimes became non-significant in the other, and the sign of the regression coefficient could also change. This finding underscores that, in practical applications, careful consideration of variable assignment is necessary for robust inference and motivates further investigation into criteria for selecting dependent and independent variables in PGLS.

However, in a more comprehensive study of the correlation between bacterial growth rate and CRISPR spacer content in genomes (Liu *et al*., 2023) using Pagel’s *λ* model (Freckleton *et al*., 2002), we found that switching the dependent and independent variables could substantially affect the results: correlations that were significant (*p* < 0.05) in one configuration sometimes became non-significant (*p* ≥ 0.05) in the other. These observations underscore that, in practical applications, careful consideration of variable assignment is necessary for robust inference and motivate further investigation into criteria for selecting dependent and independent variables in PGLS.

Using simulated data, we further assessed the frequency of inconsistent outcomes caused by switching independent and dependent variables in the PGLS analysis. In addition, we compared seven criteria to determine the appropriate designation of independent and dependent variables in the PGLS analysis using Pagel’s *λ* model (Freckleton *et al*., 2002).

## 2 MATERIALS AND METHODS

### 2.1 Empirical datasets

A dataset including the empirical minimal doubling times, CRISPR spacer numbers, optimal growth temperature, and prophage numbers of 262 bacteria was extracted from the Supplemental Material Table S1 of Liu et al. (2023). The phylogenetic tree of these 262 bacteria was retrieved from the Genome Taxonomy Database (GTDB; accessed April 8, 2022) (Parks *et al*., 2022).

### 2.2 Simulation data

First, we generated a binary tree containing 100 terminal nodes using the R package ape (Paradis & Schliep 2019). The trait *X*_1_ evolved according to a Brownian motion model (using the R package ape) with a variance rate σ^2^_*BM*_ = 4 along the tree, and then the trait *X*_2_ was simulated according to

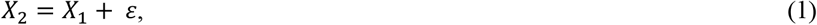

where *ε* was a normally distributed random noise term with a mean of 0 and a variance of 10^−4^, 0.01, 1, 4, 16, 64, 256 to 1024. The term *ε* introduced noise to the dependent variable *X*_2_ and a gradient variance of this noise term (from 10^−4^ to 1024) changed the correlation between *X*_1_ and *X*_2_, from strong to weak. We named this case “BM & BM + Norm.”

Then, we simulated the trait *X*_1_ according to a normal distribution with a mean of 0 and a variance of 4. The trait *X*_2_ was simulated according to equation (1), where *ε* was simulated according to a Brownian motion model with σ^2^_*BM*_ in a range of 10^−4^, 0.01, 1, 4, 16, 64, 256, 1024. We named this case “Norm & Norm + BM.”

We ran 1000 simulations for each parameter condition, and a total of 16,000 simulations were performed.

### 2.3 Software and packages

All PGLS regressions in this study were performed using the R (version 4.0.2) package phylolm (version 2.6.2) (Ho & Ane 2014). The estimated 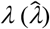, log-likelihood (LLK) (Freckleton *et al*., 2002), Akaike information criterion (AIC), *R*^2^ of the models, and the *p*-value of the regression coefficients were calculated in the PGLS regression analyses.

The phylogenetic signals (*λ* and *K*) were estimated using the R (version 4.0.2) package phytools (version 1.0-3) (Revell 2012).

## 3 RESULTS

### 3.1 Switching dependent and independent variables in PGLS analyses of empirical data sometimes gave conflicting outcomes

Using a dataset comprising a range of traits from 262 bacteria in a recent publication (Liu *et al*., 2023), we analyzed the correlations of empirical minimal doubling time and optimal growth temperature with genomic characteristics. Thirty-eight pairs of traits were re-examined using Pagel’s *λ* model in the PGLS approach. However, switching dependent and independent variables in 10 pairs resulted in conflicting outcomes, leading to different conclusions (Table 1). For instance, when considering the average prophage number as the dependent variable and the optimal growth temperature as the independent variable, the PGLS analysis showed a significant negative correlation between the two traits (*p* = 9 × 10^−4^, Table 1). In contrast, switching dependent and independent variables showed no statistically significant correlation (*p* = 0.242, Table 1). Identifying accurate findings rather than conflicting results is necessary. However, the question is how one can determine which one is correct.

**Table 1.** Switching dependent and independent variables in PGLS may yield conflicting results.^†^.

| Dependent Variable ~ Independent Variable | Slope | *p* | *R*<sup>2</sup> |
| --- | --- | --- | --- |
| EMDT ~ min_CRISPR_spacer | 0.010 | 0.049 | 0.015 |
| min_CRISPR_spacer ~ EMDT | 1.376 | 0.053 | 0.014 |
| EMDT ~ mean_prophage_number | 0.565 | 0.013 | 0.024 |
| mean_prophage_number ~ EMDT | 0.008 | 0.558 | 0.001 |
| EMDT ~ min_prophage_number | 0.622 | 0.007 | 0.028 |
| min_prophage_number ~ EMDT | 0.011 | 0.427 | 0.002 |
| EMDT ~ max_prophage_number | 0.448 | 0.037 | 0.017 |
| max_prophage_number ~ EMDT | 0.005 | 0.722 | 5×10<sup>-4</sup> |
| OPT ~ min_CRISPR_array | 0.308 | 0.061 | 0.014 |
| min_CRISPR_array ~ OPT | 0.053 | 8×10<sup>-5</sup> | 0.058 |
| OPT ~ mean_prophage_number | -0.197 | 0.242 | 0.005 |
| mean_prophage_number ~ OPT | -0.031 | 9×10<sup>-4</sup> | 0.042 |
| OPT ~ min_prophage_number | -0.154 | 0.374 | 0.003 |
| min_prophage_number ~ OPT | -0.026 | 0.004 | 0.031 |
| OPT ~ max_prophage_number | -0.183 | 0.214 | 0.006 |
| max_prophage_number ~ OPT | -0.036 | 4×10<sup>-4</sup> | 0.047 |
| OPT ~ prophage_existence | -1.046 | 0.182 | 0.007 |
| prophage_existence ~ OPT | -0.009 | 8×10<sup>-4</sup> | 0.043 |
| OPT ~ Average_Cas_gene_number | 0.197 | 0.610 | 0.002 |
| Average_Cas_gene_number ~ OPT | 0.023 | 0.012 | 0.043 |

<sup>†</sup>Pagel's $\lambda$ model was used in these analyses. The dataset containing 262 bacteria was extracted from the Supplemental Material Table S1 of Liu et al. (2023), and the phylogenetic tree used in PGLS analysis was retrieved from the Genome Taxonomy Database (Parks *et al.*, 2022). EMDT, empirical\_minimal\_doubling\_times; OPT, mean\_optimal\_growth\_temperature. The variables before and after “~” are dependent and independent.

### 3.2 Prevalence of conflicts resulting from switching dependent and independent variables

In the above survey on correlations within empirical data, we found that switching dependent and independent variables resulted in conflicting outcomes in 26.3% of cases. To evaluate the prevalence of conflicting results from switching dependent and independent variables, we simulated the progression of two traits along a binary tree of 100 species. Different distributions of the traits were simulated. In the case of “BM & BM + Norm”, we constructed the trait *X*_1_ under the Brownian motion model (“BM”) and the trait *X*_2_, as “BM + Norm”, equals to *X*_1_ plus a noise term “Norm.” In the case of “Norm & Norm + BM,” the noise term is “BM.” To account for varying levels of correlation, we set a gradient of variance in the noise term, with the variance σ^2^_*Norm*_ varying from 10^−4^, 0.01, 1, 4, 16, 64, 256, to 1024. For each parameter condition, we simulated 1000 times.

For the data from each simulation, we performed two rounds of PGLS analysis, *X*_1_∼*X*_2_ and *X*_2_∼*X*_1_ (representing regressions of *X*_1_ on *X*_2_ and *X*_2_ on *X*_1_, respectively). The results are presented in Table S1 and summarized in Table 2. In 3781 simulations, neither *X*_1_∼*X*_2_ nor *X*_2_∼*X*_1_.gave significant correlations (*p* ≥ 0.05 for all of them). In 10161 simulations, both *X*_1_∼*X*_2_ and *X*_2_∼*X*_1_.gave significant correlations (*p* < 0.05). In each of these 10161 cases, the regression coefficients of *X*_1_∼*X*_2_ and *X*_2_∼*X*_1_ have the same sign. In the other 2058 simulations, only one of *X*_1_∼*X*_2_ and *X*_2_∼*X*_1_.gave significant correlations (*p* < 0.05 for one and *p* ≥ 0.05 for the other). The frequency of conflicting results caused by switching dependent and independent variables in our simulated dataset is 12.9%.

**Table 2.** Frequency of conflicting results caused by switching dependent and independent variables in PGLS analyses using Pagel’s *λ* model.

| $X_1^\dagger$ | BM | | | | | | | | Norm | | | | | | | |
| --- | --- | --- | --- | --- | --- | --- | --- | --- | --- | --- | --- | --- | --- | --- | --- | --- |
| $X_2^\ddagger$ | BM + Norm | | | | | | | | Norm + BM | | | | | | | |
| Noise Term Variance | $10^{-4}$ | 0.01 | 1 | 4 | 16 | 64 | 256 | 1024 | $10^{-4}$ | 0.01 | 1 | 4 | 16 | 64 | 256 | 1024 |
| Simulation times | $10^3$ | $10^3$ | $10^3$ | $10^3$ | $10^3$ | $10^3$ | $10^3$ | $10^3$ | $10^3$ | $10^3$ | $10^3$ | $10^3$ | $10^3$ | $10^3$ | $10^3$ | $10^3$ |
| $X_1 \sim X_2$ gave $p < 0.05^\S$ | $10^3$ | $10^3$ | $10^3$ | 982 | 543 | 188 | 68 | 53 | $10^3$ | $10^3$ | $10^3$ | $10^3$ | 926 | 387 | 134 | 74 |
| $X_2 \sim X_1$ gave $p < 0.05$ | $10^3$ | $10^3$ | $10^3$ | $10^3$ | 954 | 528 | 171 | 81 | $10^3$ | $10^3$ | $10^3$ | $10^3$ | $10^3$ | 844 | 318 | 129 |
| Both $X_1 \sim X_2$ and $X_2 \sim X_1$ gave $p < 0.05$ | $10^3$ | $10^3$ | $10^3$ | 982 | 534 | 169 | 34 | 14 | $10^3$ | $10^3$ | $10^3$ | $10^3$ | 926 | 375 | 88 | 39 |
| Both $X_1 \sim X_2$ and $X_2 \sim X_1$ gave $p \geq 0.05$ | 0 | 0 | 0 | 0 | 37 | 453 | 795 | 880 | 0 | 0 | 0 | 0 | 0 | 144 | 636 | 836 |
| $X_1 \sim X_2$ and $X_2 \sim X_1$ gave conflicting results<sup>¶</sup> | 0 | 0 | 0 | 18 | 429 | 378 | 171 | 106 | 0 | 0 | 0 | 0 | 74 | 481 | 276 | 125 |

<sup>†</sup>The trait $X_1$ evolved according to a Brownian motion (BM) model in the first 8000 simulations, and evolved according to a normal distribution in the other 8000 simulations. For details, see the MATERIALS AND METHODS section.
<sup>‡</sup>The trait $X_2$ was simulated by adding a noise term to $X_1$ . The term noise was normally distributed in the first 8000 simulations and generated according to a Brownian motion model in the other 8000 simulations. For details, see the MATERIALS AND METHODS section.
<sup>§</sup>The variables before and after “ ∼ ” are dependent and independent, respectively.
<sup>¶</sup>When one of $X_1 \sim X_2$ and $X_2 \sim X_1$ gives a $p < 0.05$ result and the other gives a $p \geq 0.05$ result, we define these results as conflicting.

Table 2 shows a relationship between the variance in simulations and the frequency of conflicting results caused by switching dependent and independent variables. When the variance of the noise term is slight (like 10^−4^, 0.01, 1, and 4), i.e., where there is a strong correlation between *X*_1_ and *X*_2_, switching independent and dependent variables gives almost the same results. As the variance in the term noise increases (i.e., where the correlation between *X*_1_ and *X*_2_ becomes weak), switching independent and dependent variables leads to many conflicting results. In cases where the variance is 16 in “BM & BM + Norm” and 64 in “Norm & Norm + BM,” there are even close to 50% of cases with conflicting results. However, as the variance becomes much more pronounced and the correlation between *X*_1_ and *X*_2_ becomes much weaker, the frequency of conflicting results caused by switching dependent and independent variables decreases (Table 2).

### 3.3 Estimated lambda may change according to the choice of the dependent variable

To control the phylogenetic dependence in the data, PGLS assumes that the model residual error is distributed according to the variance-covariance matrix representing the residual variance and the evolutionary relationships among species. In its simplest form, PGLS assumes a Brownian motion model of evolution at a single rate. By parameter estimation and altering the properties of the variance-covariance matrix, PGLS can be extended to various evolutionary models, like Pagel’s *λ* model (Freckleton *et al*., 2002), Ornstein–Uhlenbeck model (Hansen 1997), and the early burst model (Harmon *et al*., 2010). We suspect that switching independent and dependent variables in the PGLS analysis may result in inconsistent estimates of the parameters (like Pagel’s *λ*) and so inconsistent correction for the phylogenetic non-independence of the trait values.

For the data of each simulation, we calculated the phylogenetic signal *λ* of the two traits (*X*_1_ and *X*_2_) and the 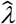 of *X*_1_∼*X*_2_ and *X*_2_∼*X*_1_ using Pagel’s *λ* model.

In the “BM & BM + Norm” case, when the variance of the noise term is slight, both *X*_1_ and *X*_2_ exhibit strong phylogenetic signals with their *λ* values close to 1, and both *X*_1_∼*X*_2_ and *X*_2_∼*X*_1_ give small 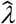 values (Figure 1). However, with the increase of the variance of the noise term, the *λ* values of *X*_2_ gradually decrease while those of *X*_1_. remain at their high levels. Together, the 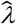 values of the *X*_1_∼*X*_2_ models gradually increase to 1 while those of the *X*_2_∼*X*_1_ models remain stable at around 0.03 (Figure 1).

**Figure 1.**
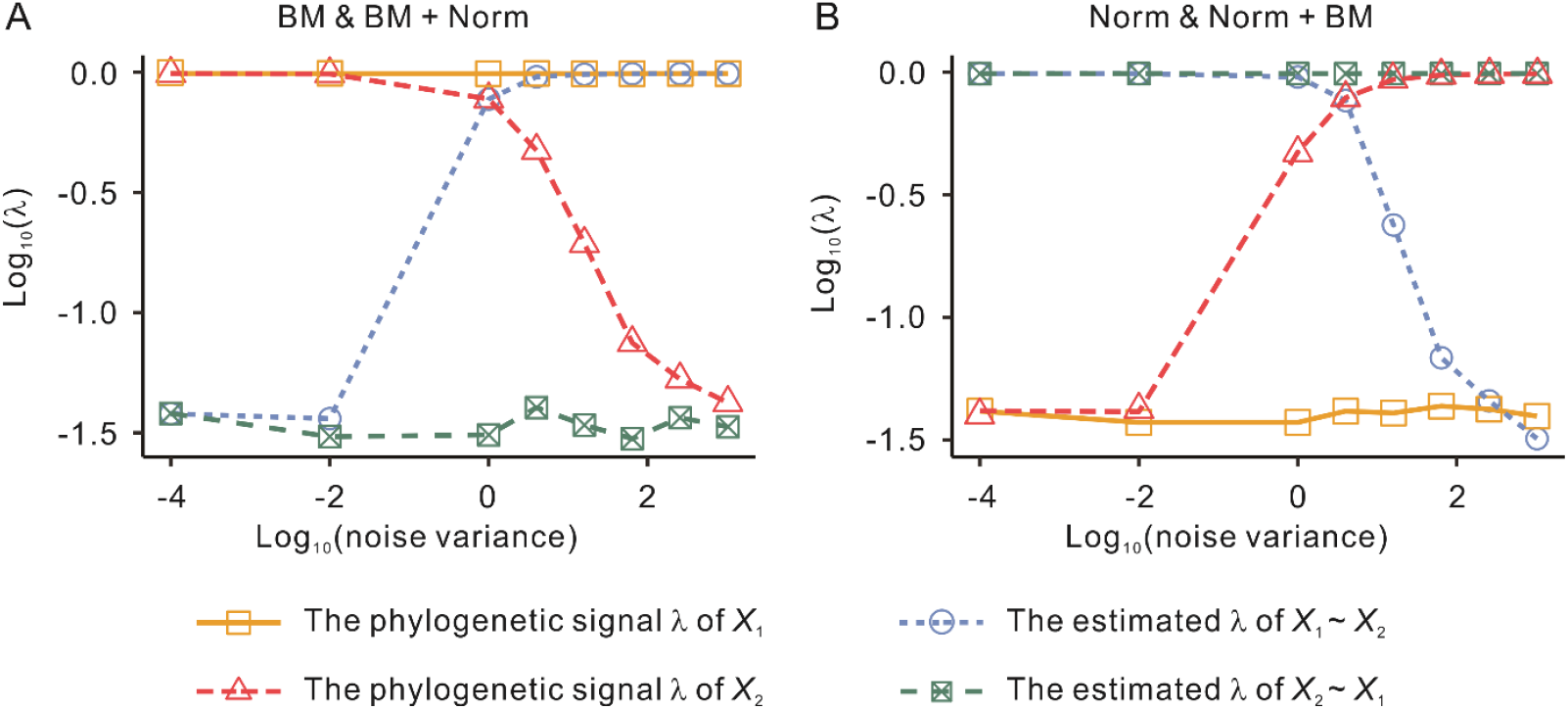
Phylogenetic signals of the variables and the estimated *λ* of *X*_1_∼*X*_2_ and *X*_2_∼*X*_1_ models under different noise term variances. For each variance gradient, 1000 simulations were conducted. The data points on the graph represent the average value of phylogenetic diagnostics (phylogenetic signals or model estimated *λ*) of the noise term under a specific variance gradient based on the 1000 simulations. (A) In the “BM & BM + Norm” case, *X*_1_ was simulated according to a Brownian motion (BM) model, while *X*_2_ was simulated based on *X*_1_ plus a normally distributed random noise term. (B) In the “Norm & Norm + BM” case, *X*_1_ was simulated according to a normal distribution, and *X*_2_ was simulated according to *X*_1_ plus a noise term, where the noise term followed a BM model. *X*_1_∼*X*_2_ and *X*_2_∼*X*_1_ represent PGLS regressions using *X*_1_ and *X*_2_ as dependent variables, respectively.

In the “Norm & Norm + BM” case, we found a pattern previously reported by Revell (2010), PGLS regression is possible to give a high 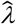 value even if both traits have very weak phylogenetic signals. When the variance of the BM noise term is slight, both *X*_1_ and *X*_2_ have small *λ* values but both *X*_1_∼*X*_2_ and *X*_2_∼*X*_1_ models yielded high 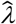 values of about 1 (Figure 1). With the increase of the variance of the noise term, the phylogenetic signals of *X*_2_ gradually become stronger and finally approach one, but that of *X*_1_ remains weak. Meanwhile, the 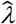 values of *X*_1_∼*X*_2_ gradually decrease to about 0.03, while those of *X*_2_∼*X*_1_ remain close to 1 (Figure 1).

We could see that the two models, *X*_1_∼*X*_2_ and *X*_2_∼*X*_1_, generally give similar 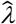 values when the two variables, *X*_1_ and *X*_2_, have similar *λ* values (Figure 1). That is, switching dependent and independent variables would not dramatically affect the parameter estimation of the PGLS regression analyses when the two analyzed traits have similar levels of phylogenetic signals. However, when the two variables differ strikingly in their phylogenetic signals, the two models give quite different 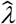 values (Figure 1). In this case, switching dependent and independent variables would dramatically affect the parameter estimation and potentially give conflicting results. In addition, we noticed that the 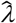 values tend to be close to the phylogenetic signal of the dependent variable.

### 3.4 A golden standard to evaluate the accuracy of PGLS results

When performing PGLS analysis on both empirical and simulated datasets, switching dependent and independent variables leads to a significant frequency of conflicting results. Avoiding arbitrary selection of the dependent variable is essential in PGLS analysis using Pagel’s *λ* model. In order to select a better dependent variable, we will use simulated data to establish a selection criterion.

In empirical phylogenetic data, we only have access to trait values at terminal nodes, and potential correlations between traits can be estimated using methods such as PGLS. In contrast, we can access trait values at internal nodes in simulated phylogenetic data. Since changes along different phylogenetic branches are independent, we can measure the evolutionary correlation between two traits by analyzing their changes along phylogenetic branches using standard statistical methods. First of all, we calculated the changes in the traits *X*_1_ and *X*_2_ along evolutionary branches per unit time, Δ*X*_1_/*L* and Δ*X*_2_/*L*, where *L* is the length of the branch. Then, we did Shapiro-Wilk test on Δ*X*_1_/*L* and Δ*X*_2_/*L* to see if they had normal distributions. If both of them satisfy normality, we used Pearson correlation to detect the correlation between Δ*X*_1_/*L* and Δ*X*_2_/*L*. Otherwise, we used Spearman’s rank correlation. These analyses revealed significant correlations between the two traits, *X*_1_ and *X*_2_, in 11911 simulations (10988 positives and 923 negatives, *p* < 0.05 for all these cases) but not in the other 4089 simulations (*p* ≥ 0.05 for all these cases) (Table S1). These results provide “golden standards” for judging whether PGLS analyses of trait values on terminal nodes (*X*_1_ and *X*_2_) give correct results.

PGLS analyses of *X*_1_∼*X*_2_ show that there are significant positive correlations in 10326 simulations (*p* < 0.05 for all these cases), significant negative correlations in 29 simulations (*p* < 0.05 for all these cases), and no significant correlations in 5645 simulations (*p* ≥ 0.05 for all these cases) (Table S1). The same analyses of *X*_2_∼*X*_1_ show that there are significant positive correlations in 12005 simulations (*p* < 0.05 for all these cases), significant negative correlations in 20 simulations (*p* < 0.05 for all these cases), and no significant correlations in 3975 simulations (*p* ≥ 0.05 for all these cases) (Table S1). By comparing with the golden standards, we found that PGLS analyses gave correct results for both *X*_1_∼*X*_2_ and *X*_2_∼*X*_1_ in 11474 simulations (71.71%) and incorrect results for both *X*_1_∼*X*_2_ and *X*_2_∼*X*_1_ in 2468 simulations (15.43%). In 2058 (12.86%) simulations, only one of the two competing models, *X*_1_∼*X*_2_ or *X*_2_∼*X*_1_, yielded correct results. Therefore, limited by the performance of PGLS regression analysis, we could at most obtain an accuracy of 84.57% in analyzing the data of our 16000 simulations by PGLS regression. However, if we arbitrarily select one trait (*X*_1_ or *X*_2_) as the independent variable with the worst luck, we could get an accuracy of only 71.71%. In the following attempt to find an accurate criterion for the selection of dependent variables, we hope to perceive more correct cases from the 2058 simulations, where *X*_1_∼*X*_2_ and *X*_2_∼*X*_1_ gave conflicting results.

### 3.5 Looking for an accurate criterion for dependent variable selection

Referring to the golden standards, we evaluated the potential performance of seven criteria for selecting a better model from *X*_1_∼*X*_2_ and *X*_2_∼*X*_1_.

In statistics, the suitability of two competing statistical models is often assessed by calculating each model’s log-likelihood (LLK) value. First, we examined whether a higher (or lower) LLK could give an accurate prediction of the better model between *X*_1_∼*X*_2_ and *X*_2_∼*X*_1_. By calculating the LLKs of the two models for the 2058 simulations (Table S1), we found that the models selected from *X*_1_∼*X*_2_ and *X*_2_∼*X*_1_ by a lower LLK (denoted as *Model*_*LLK,lower*_) have more correct outcomes than the alternative model (denoted as *Model*_*LLK,higher*_), 1076 *vs*. 982. A *χ*^2^ test showed a statistically significant difference (*p* = 3.7 × 10^−3^, Table 3).

**Table 3.** Differences in the performance of models selected by seven criteria.

| Criteria | Models | Correct | Incorrect | $\chi^2$ | $p^\dagger$ |
| --- | --- | --- | --- | --- | --- |
| LLK | $Model_{LLK,higher}$ | 982 | 1076 | 8.405 | $3.7 \times 10^{-3}$ |
| | $Model_{LLK,lower}$ | 1076 | 982 | | |
| AIC | $Model_{AIC,higher}$ | 1076 | 982 | 8.405 | $3.7 \times 10^{-3}$ |
| | $Model_{AIC,lower}$ | 982 | 1076 | | |
| $R^2$ | $Model_{R^2,higher}$ | 1088 | 970 | 13.30 | $2.7 \times 10^{-4}$ |
| | $Model_{R^2,lower}$ | 970 | 1088 | | |
| $p$ | $Model_{p,higher}$ | 970 | 1088 | 13.30 | $2.7 \times 10^{-4}$ |
| | $Model_{p,lower}$ | 1088 | 970 | | |
| $\lambda_y$ | $Model_{\lambda_y,higher}$ | 1734 | 324 | 1929 | $< 2.2 \times 10^{-16}$ |
| | $Model_{\lambda_y,lower}$ | 324 | 1734 | | |
| $K_y$ | $Model_{K_y,higher}$ | 1734 | 324 | 1929 | $< 2.2 \times 10^{-16}$ |
| | $Model_{K_y,lower}$ | 324 | 1734 | | |
| $\hat{\lambda}$ | $Model_{\hat{\lambda},higher}$ | 1734 | 324 | 1929 | $< 2.2 \times 10^{-16}$ |
| | $Model_{\hat{\lambda},lower}$ | 324 | 1734 | | |

<sup>†</sup>The minimal significance value that the R environment (version 4.1.0) gives is $2.2 \times 10^{-16}$ .

Akaike information criterion (AIC) is a widely used estimator of the quality of statistical models for a given dataset (Akaike 1974). It balances the excellent fit of the model with the complexity of the model. Calculating the AIC values of the two competing models for the 2058 simulations (Table S1), we found that the models selected with a higher AIC value (denoted as *Model*_*AIC,higher*_) had significantly more correct outcomes than the alternative model (denoted as *Model*_*AIC,lower*_), 1076 *vs*. 982, *p* = 3.7 × 10^−3^ (Table 3).

*R*^2^ describes the proportion of total variation in the dependent variable explained by independent variables in the regression model, and the *p*-value of the regression coefficient is the probability of observing the test statistic value under the assumption that the null hypothesis is true, where the regression coefficient equals 0. These two parameters are widely used indicators of the suitability of regression models. The models selected by a higher *R*^2^ (denoted as 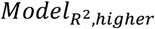) and a lower *p*-value (denoted as *Model*_*p,lower*_) have significantly more correct outcomes than the alternative models (*p* = 2.7 × 10^−4^ for both cases, Table 3).

The phylogenetic signal measures how much phylogenetic structure influences species trait values. Pagel’s *λ* and Blomberg et al.’s *K* are the most commonly used indicators of the phylogenetic signal (Pagel 1999; Blomberg *et al*., 2003). The estimated 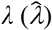 is a parameter in Pagel’s *λ* model (Pagel 1997) that measures the relatedness of the regression residuals with the phylogenetic structure. We found that the models that use the trait with a higher *λ* value or a higher *K* value as the dependent variable (denoted as 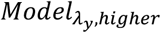 and 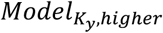, respectively) have significantly more correct outcomes than the alternate models (denoted as 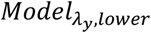 and 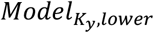, respectively) (*p* < 2.2 × 10^−16^ for both case Table 3). And the model chosen by a higher 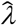 (denoted as 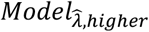) also had significantly more correct results than the alternative model (denoted as 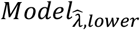) (*p* < 2.2 × 10^−16^, Table 3).

In addition, we defined a virtual criterion where one model was randomly selected from the two competing models, *X*_1_∼*X*_2_ and *X*_2_∼*X*_1_. The results of the models selected by this virtual criterion (*Model*_*rc*_) were compared with the better models selected by the above seven criteria. Although the better models selected by the above seven criteria consistently have more correct results than *Model*_*rc*_ (Table 4), none of *Model*_*LLK,lower*_, *Model*_*AIC,higher*_, 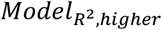, and *Model*_*p,lower*_ is significantly different from *Model*_*rc*_ (*p* > 0.100 for all four cases), but the differences of 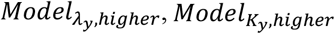, and 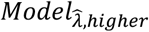 with *Model*_*rc*_ are highly significant (*p* < 2.2 × 10^−16^ for all the three cases).

**Table 4.**
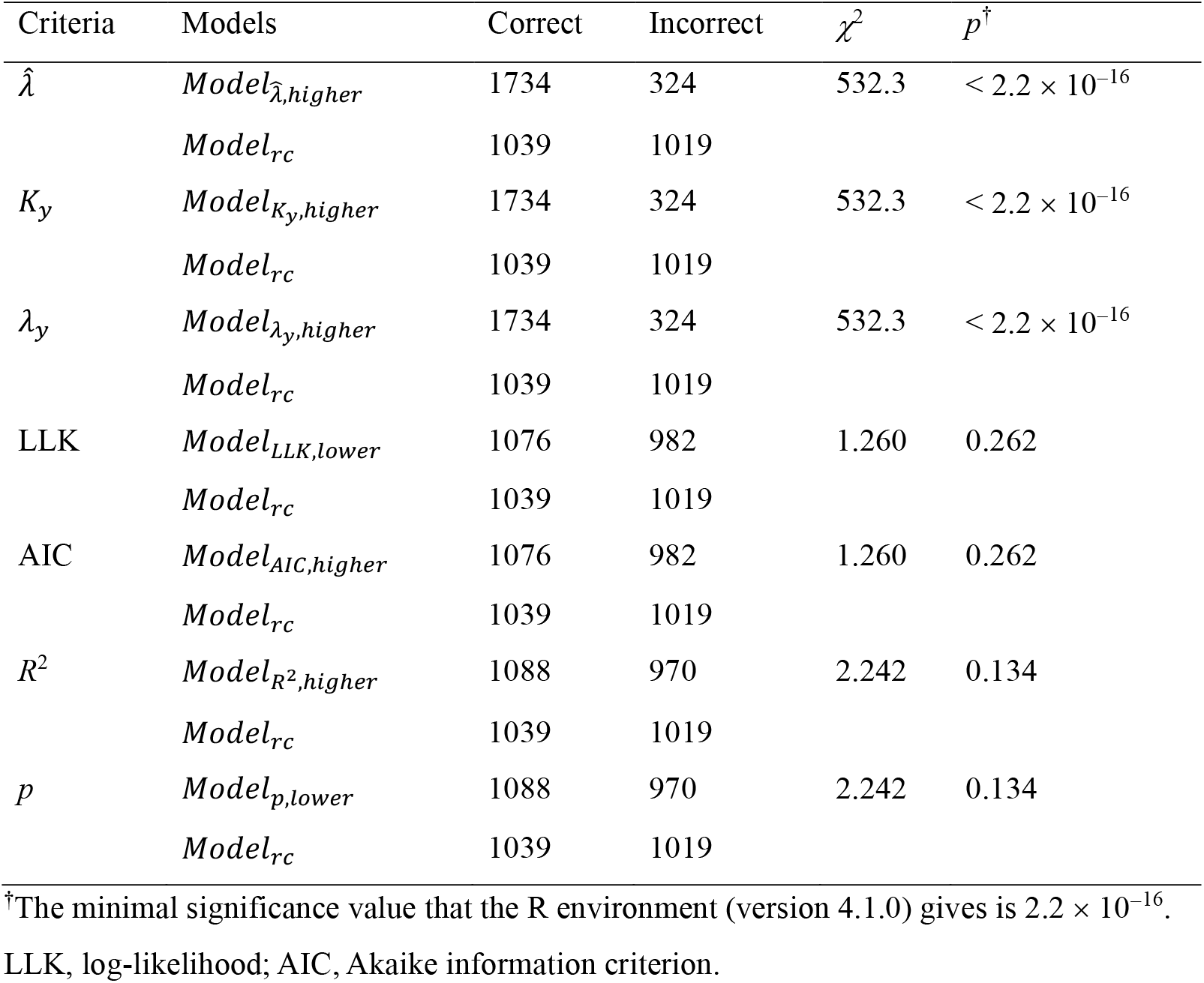
Comparing the performance of seven criteria with random selection.

| Criteria | Models | Correct | Incorrect | $\chi^2$ | $p^\dagger$ |
| --- | --- | --- | --- | --- | --- |
| $\hat{\lambda}$ | $Model_{\hat{\lambda},higher}$ | 1734 | 324 | 532.3 | $< 2.2 \times 10^{-16}$ |
| | $Model_{rc}$ | 1039 | 1019 | | |
| $K_y$ | $Model_{K_y,higher}$ | 1734 | 324 | 532.3 | $< 2.2 \times 10^{-16}$ |
| | $Model_{rc}$ | 1039 | 1019 | | |
| $\lambda_y$ | $Model_{\lambda_y,higher}$ | 1734 | 324 | 532.3 | $< 2.2 \times 10^{-16}$ |
| | $Model_{rc}$ | 1039 | 1019 | | |
| LLK | $Model_{LLK,lower}$ | 1076 | 982 | 1.260 | 0.262 |
| | $Model_{rc}$ | 1039 | 1019 | | |
| AIC | $Model_{AIC,higher}$ | 1076 | 982 | 1.260 | 0.262 |
| | $Model_{rc}$ | 1039 | 1019 | | |
| $R^2$ | $Model_{R^2,higher}$ | 1088 | 970 | 2.242 | 0.134 |
| | $Model_{rc}$ | 1039 | 1019 | | |
| $p$ | $Model_{p,lower}$ | 1088 | 970 | 2.242 | 0.134 |
| | $Model_{rc}$ | 1039 | 1019 | | |

<sup>†</sup>The minimal significance value that the R environment (version 4.1.0) gives is $2.2 \times 10^{-16}$ .
LLK, log-likelihood; AIC, Akaike information criterion.

From Tables 3-4 and S2, we can see three distinct groups of criteria: LLK and AIC, *R*^2^ and *p*-value, and Pagel’s *λ*, Blomberg et al.’s *K*, and 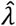. Within each group, the criteria gave the same results. Among the three groups, Pagel’s *λ*, Blomberg et al.’s *K*, and 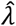 seem to be the best criterion for dependent variable selection. For a quantitative evaluation of these impressions, we performed the *χ*^2^ tests to compare Pagel’s λ with Blomberg et al.’s *K*, 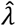, LLK, AIC, *R*^2^, and *p*-value using the 2058 simulations where *X*_1_∼*X*_2_ and *X*_2_∼*X*_1_ gave conflicting results. As shown in Table 5, the equivalency among Pagel’s *λ*, Blomberg et al.’s *K*, and 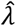 and the superiority of *λ* to LLK, AIC, *R*^2^, and *p*-value have been statistically confirmed.

**Table 5.** Comparing the performance of *λ*_*y*_ with other criteria for dependent variable selection.

| Criteria | Models | Correct | Incorrect | $\chi^2$ | $p^\dagger$ |
| --- | --- | --- | --- | --- | --- |
| $\lambda_y$ vs. $\hat{\lambda}$ | $Model_{\lambda_y, higher}$ | 1734 | 324 | 0 | 1 |
| | $Model_{\hat{\lambda}, higher}$ | 1734 | 324 | | |
| $\lambda_y$ vs. $K_y$ | $Model_{\lambda_y, higher}$ | 1734 | 324 | 0 | 1 |
| | $Model_{K_y, higher}$ | 1734 | 324 | | |
| $\lambda_y$ vs. LLK | $Model_{\lambda_y, higher}$ | 1734 | 324 | 484.1 | $< 2.2 \times 10^{-16}$ |
| | $Model_{LLK, lower}$ | 1076 | 982 | | |
| $\lambda_y$ vs. AIC | $Model_{\lambda_y, higher}$ | 1734 | 324 | 484.1 | $< 2.2 \times 10^{-16}$ |
| | $Model_{AIC, higher}$ | 1076 | 982 | | |
| $\lambda_y$ vs. $R^2$ | $Model_{\lambda_y, higher}$ | 1734 | 324 | 468.9 | $< 2.2 \times 10^{-16}$ |
| | $Model_{R^2, higher}$ | 1088 | 970 | | |
| $\lambda_y$ vs. $p$ | $Model_{\lambda_y, higher}$ | 1734 | 324 | 468.9 | $< 2.2 \times 10^{-16}$ |
| | $Model_{p, lower}$ | 1088 | 970 | | |

<sup>†</sup>The minimal significance value that the R environment (version 4.1.0) gives is $2.2 \times 10^{-16}$ . LLK, log-likelihood; AIC, Akaike information criterion.

In summary, Pagel’s *λ*, Blomberg et al.’s *K*, and 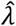 are the best criteria for dependent variable selection. Of the 2058 simulations that *X*_1_∼*X*_2_ and *X*_2_∼*X*_1_ gave conflicting results, these criteria led to correct results in 1734 simulations. Combined with the 11474 simulations where both *X*_1_∼*X*_2_ and *X*_2_∼*X*_1_ gave correct results, PGLS analysis was able to achieve 82.55% accuracy when the trait with a stronger phylogenetic signal was selected as the dependent variable.

## 4 DISCUSSION

In the PGLS correlation analysis of an empirical dataset (Liu *et al*., 2023), we found that switching dependent and independent variables could lead to a significant incidence of conflicting results. To investigate this further, we simulated the evolution of two traits (*X*_1_ and *X*_2_) along a binary tree with 100 terminal nodes, using different models and variances for 16000 iterations. Our PGLS analyses of these simulated data showed that the frequency of conflicting results caused by switching dependent and independent variables depends on the strength of the relationship between the analyzed traits. Furthermore, we showed that switching dependent and independent variables leads to different parameter estimations in PGLS regression analysis of two traits that have quite different phylogenetic signals. Using simulation, we established a golden standard for whether the two traits, *X*_1_ and *X*_2_, are correlated in each simulation by conventional correlation analysis of the changes of the two traits along the branches of the phylogenetic tree. With this golden standard, we can tell which model, *X*_1_∼*X*_2_ or *X*_2_∼*X*_1_, is correct. Seven potential criteria for dependent variable selection, LLK, AIC, *R*^2^, *p*-value, Pagel’s *λ*, Blomberg et al.’s *K*, and 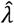, have been compared. The last three criteria are equivalent in selecting dependent variables and have shown their superiority over the other four criteria. Pagel’s *λ* and Blomberg et al.’s *K* values are generally calculated at the beginning of the phylogenetic comparative analysis, so they are already known before the PGLS analysis. If we can choose the trait with a higher *λ* or *K* value as the dependent variable, two rounds of PGLS analysis, like the *X*_1_∼*X*_2_ and *X*_2_∼*X*_1_, are not required. Considering the practical convenience, Pagel’s *λ* and Blomberg et al.’s *K* are superior to 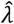.

As suggested by some previous publications, like Cooper et al. (2016), Paterno et al. (2018), and Uyeda et al. (2018), PGLS regression is not a perfect method for phylogenetic correlation analysis. In our 16000 simulations, PGLS regression analyses yielded incorrect results for 2468 cases, regardless of which trait is designated as the dependent variable. In 2058 simulations, the two competing models, *X*_1_∼*X*_2_ or *X*_2_∼*X*_1_, gave conflicting results: one is right and the other is wrong. That is, PGLS regression analyses of 16000 simulations with arbitrarily selected dependent variables could yield correct results in the range of 71.79% to 84.57%. By selecting the trait with a strong phylogenetic signal as the dependent variable, we could obtain an accuracy of 82.55%. As this accuracy is still 2.02% below the upper limit of PGLS regression using Pagel’s *λ* model, a better criterion may be found in the future. PGLS regression analysis itself should also be improved.

It is important to emphasize that the terms “independent” and “dependent” variables can be misleading when using PGLS to study the correlation between traits whose causal relationships are unclear, and should not be interpreted literally. In such cases, the independent variable does not necessarily represent the cause of changes in the dependent variable. PGLS is thus primarily designed to account for phylogenetic non-independence and effectively replaces conventional correlation methods, such as Pearson and Spearman’s rank tests, in comparative studies. When selecting a dependent variable for PGLS, the focus should be on obtaining robust and interpretable estimates of the relationship rather than assigning causal roles. Specifically, even if a trait appears to drive changes in another biologically, it should be designated as the dependent variable if it exhibits a stronger phylogenetic signal, as this choice generally produces more consistent results.

This study specifically addresses the issue of dependent variable selection in PGLS analyses using Pagel’s λ model (Freckleton *et al*., 2002). To examine the theoretical consistency, we performed PGLS analyses on 16,000 simulated datasets under a Brownian motion model, alternating *X*_1_ and *X*_2_ as the dependent variable (Table S1). For all simulations, the results of *X*_1_∼*X*_2_ and *X*_2_∼*X*_1_ were consistent in terms of significance (*p* value) and the sign of the regression coefficient, indicating that arbitrary choice of the dependent variable does not affect correlation estimates in models without parameter estimation. In contrast, for models involving parameter estimation, such as the Ornstein–Uhlenbeck model (Hansen 1997) or the early burst model (Harmon *et al*., 2010), swapping dependent and independent variables can produce different parameter estimates and potentially conflicting conclusions. Further research is needed to establish clear criteria for selecting dependent variables in these models.

Separately, in a more comprehensive empirical study of the correlation between bacterial growth rate and CRISPR spacer content in genomes (Liu et al., 2023), we observed that switching dependent and independent variables could change the significance of the correlation. This observation highlights that in practical applications, especially with real biological data, careful consideration of variable assignment—favoring traits with stronger phylogenetic signal—is critical for robust and reproducible inference in PGLS.

## ACKNOWLEDGMENTS

This work was supported by the National Natural Science Foundation of China (Grant number 31671321).

## DATA AVAILABILITY STATEMENT

Supplementary Tables S1-S2 are submitted as Supporting Information for Online Publication.

## AUTHOR CONTRIBUTIONS

Deng-Ke Niu and Zheng-Lin Chen conceived the idea and designed the general methodology; Zheng-Lin Chen designed the simulations; Zheng-Lin Chen and Hong-Ji Guo implemented the simulations and data analysis; Deng-Ke Niu led the writing of the manuscript. All authors contributed critically to the drafts and gave final approval for publication.

