## Supplemental Table S2 for "Dependent variable selection in phylogenetic generalized least squares regression analysis under Pagel’s lambda model"

Table S1. Differences in the performance of models selected by six criteria in 12000 simulations.

| Criteria | Models | Correct | Incorrect | *χ*^2^ | *p*^a^ |
| --- | --- | --- | --- | --- | --- |
| LLK | ${Model}_{LLK,higher}$ | 12456 | 3544 | 1.583 | 0.208 |
|  | ${Model}_{LLK,lower}$ | 12550 | 3450 |  |  |
| AIC | ${Model}_{AIC,higher}$ | 12550 | 3450 | 1.583 | 0.208 |
|  | ${Model}_{AIC,lower}$ | 12456 | 3544 |  |  |
| *R*^2^ | ${Model}_{R^{2},higher}$ | 12562 | 3438 | 2.505 | 0.114 |
|  | ${Model}_{R^{2},lower}$ | 12444 | 3556 |  |  |
| *p* | ${Model}_{p,higher}$ | 12444 | 3556 | 2.505 | 0.114 |
|  | ${Model}_{p,lower}$ | 12562 | 3438 |  |  |
| $\lambda_{y}$ | ${Model}_{\lambda_{y},higher}$ | 13208 | 2792 | 363.3 | < 2.2 × 10^−16^ |
|  | ${Model}_{\lambda_{y},lower}$ | 11798 | 4202 |  |  |
| $\hat{\lambda}$ | ${Model}_{\hat{\lambda},higher}$ | 13208 | 2792 | 363.3 | < 2.2 × 10^−16^ |
|  | ${Model}_{\hat{\lambda},lower}$ | 11798 | 4202 |  |  |
| $K_{y}$ | ${Model}_{K_{y},higher}$ | 13208 | 2792 | 363.3 | < 2.2 × 10^−16^ |
|  | ${Model}_{K_{y},higher}$ | 11798 | 4202 |  |  |

^a^The minimal significance value that the R environment (version 4.1.0) gives is 2.2 × 10^−16^. LLK, log-likelihood; AIC, Akaike information criterion.
